# Spatially resolved transcriptomics reveals unique gene signatures associated with human temporal cortical architecture and Alzheimer’s pathology

**DOI:** 10.1101/2021.07.07.451554

**Authors:** Shuo Chen, Yuzhou Chang, Liangping Li, Diana Acosta, Cody Morrison, Cankun Wang, Dominic Julian, Mark E. Hester, Geidy E. Serrano, Thomas G. Beach, Qin Ma, Hongjun Fu

**Author notes:** Both authors contributed equally (Shuo Chen, Yuzhou Chang). Corresponding Authors: H.F.; Q.M.

## Abstract

Alzheimer’s disease (AD) is pathologically characterized by amyloid beta (Aβ) plaques, neurofibrillary tangles (tau aggregates), and alterations in microglia, astrocytes and oligodendrocytes. The mesial temporal lobe is a vulnerable brain region in early AD; however, little is known about the transcriptome-scale gene expression in this region and its relation to AD pathology. Here we use the 10x Genomics Visium platform in combination with co-immunofluorescence staining of AD-associated pathological markers to define the spatial topography of gene expression in the middle temporal gyrus (MTG) from both early AD and age- and gender-matched control cases. We identify unique marker genes for six cortical layers and the adjacent white matter as well as gene expression patterns and alterations that showcase unique gene signatures and pathways associated with a range of AD pathology. Also, gene co-expression analyses of differentially expressed genes (DEGs) between AD and controls reveal four unique gene modules, which significantly change their co-expression patterns in the presence of variations of AD pathology. Furthermore, we validate the changes of key representative DEGs that are associated with AD pathology in neurons, microglia, astrocytes and oligodendrocytes using single-molecule fluorescent in situ hybridization. In summary, we provide a rich resource for the spatial transcriptomic profile of the human MTG, which will contribute to our understanding of the complex architecture and AD pathology of this vulnerable brain region.

## Main

Alzheimer’s disease (AD) is the most common form of dementia in the elderly, affecting 40-50 million people worldwide^1^. AD is pathologically characterized by extracellular amyloid beta (Aβ) plaques, neurofibrillary tangles (neuronal tau aggregates), gliosis induced by activated microglia and reactive astrocytes, and white matter degeneration, possibly induced by dysfunctions in oligodendrocytes^2–5^. The stereotypical spread of AD pathology from regions like the medial temporal lobe to the cortex has been extensively studied^6, 7^. However, the molecular mechanisms underlying the cell- and region-specific distribution of AD pathology at early stages of the disease are still not fully elucidated. The transcriptome of the AD brain can pinpoint key differences in disease that may be crucial for elucidating the pathogenesis of AD and for developing disease-modifying therapeutics for the prevention and treatment of AD^8^.

Compared with the single-cell or single-nucleus (sn) RNA-Sequencing (RNA-Seq) techniques, the recent advent of spatially resolved transcriptomics (ST) (e.g., 10x Genomics Visium) allows us to compare transcriptomic profiles between AD and control subjects without the loss of spatial information^9–14^. This novel technique will allow for a better understanding of molecular mechanisms underlying the neuropathology of AD within the spatial context. Recent applications of ST on AD-like mouse models have revealed a plaque-induced gene (PIG) network, an oligodendrocyte gene (OLIG) response, and regionally differentially expressed genes (DEGs) in AD-like mice compared to controls^13, 14^. However, comprehensive ST profiling of human AD brain tissue and characterization of DEGs associated with various AD pathology, have not been reported as far as we know.

To address this knowledge gap, we used the 10x Genomics Visium platform in combination with co-immunofluorescence staining of AD-associated pathological markers to define the spatial topography of gene expression in the human middle temporal gyrus (MTG), a vulnerable brain region in early AD^15^. In this study, we chose the MTG from human postmortem non-AD control (Braak stages I-II, very little tau pathology) and early AD cases (Braak stages III-IV)^7^. In order to highlight gene changes associated with early stages of disease and exclude the majority of downstream gene expression changes that occur in late AD, we did not include AD cases at Braak stages V-VI in this study. We identified specific marker genes for six cortical layers and the white matter as well as unique genes and pathways associated with various AD pathology. Weighted gene co-expression network analyses (WGCNA)^16^ of DEGs between early AD and control cases revealed four unique co-expression gene modules (M1-4), which are expressed in four main cell types (neurons, microglia, astrocytes, and oligodendrocytes). The co-expression pattern of M1-4 significantly changed in the presence of specific AD pathology. Furthermore, we quantitated the expression of key representative DEGs associated with AD pathology in neurons, microglia, astrocytes and oligodendrocytes of human early AD and control cases using RNAscope single-molecule fluorescent in situ hybridization (smFISH). The resulting resource is the first, as far as we know, to provide ST profiling of human MTG and a unique spatial view of transcriptional alterations associated with various AD pathology. Furthermore, this analysis reveals gene perturbations specific to each layer- and AD pathology, as well as shared gene-expression perturbations, thus providing additional molecular insights into AD.

## Results

### ST in human postmortem MTG from AD and control cases

Human postmortem fresh frozen brain sections from the MTG of two control (Braak stages I-II, male, 72 and 82 years old) and two early AD (Braak III-IV, male, 86 and 89 years old) cases (Supplementary Table 1) were chosen to perform the 10x Genomics Visium ST experiments. For each sample, seven continuous sections (10 µm) were saved and labeled sequentially as No.1 to No.7. The middle section (No.4) was mounted on the Visium Gene Expression (GE) slide for ST profiling. The outmost two sections (No.1 and No.7) were used for RNAscope HiPlex assay. The remaining sections (No. 2, 3, 5, 6) were used for immunofluorescence (IF) staining of cell-type markers and AD pathology. We obtained a total of 17,203 ST spots from all four samples, and we detected 8638 ± 5863 unique molecular identifiers (UMIs) and 3340 ± 1701 unique genes per ST spot (Supplementary Table 2). 250 spots were classified as noise and not used for further analysis because of tissue loss or damage or ambiguous clustering results in uniform manifold approximation and projection (UMAP).

Although the 10x Genomics Visium platform cannot attain the single-cell resolution yet, the diameter of each ST spot (55 µm) should cover 2-5 cells. Since the thickness of each brain section used in this study is 10 µm, the total distance from section No.2 to No.6 is 50 µm, which is less than the diameter of a ST spot. Given that all the adjacent sections are within 20 µm from the GE section (No.4) and the average cell-body size of most nerve cells in the human brain is ∼ 10-20 µm, the cell distribution and brain architecture can be used to align adjacent sections to the GE section^13^. Therefore, we aligned the GE section with the hematoxylin and eosin (H&E) staining and 4 adjacent sections with IF staining to annotate the cortical layers, the white matter and AD pathology-associated regions. The No.2 section was co-stained with WFS1/AT8, No.3 with P2RY12/GFAP/AT8, No.5 with Olig2/Tau46/Aβ, and No.6 with TauC/GAD1 antibodies (Fig. 1a). WFS1 (wolframin) were used as layer II marker^17^. AT8 and Aβ antibodies were used to stain pathological tau and Aβ plaques, respectively. GAD1, P2RY12, GFAP and Oligo2 were used as cell type-specific markers for inhibitory neurons, microglia, astrocytes, and oligodendrocytes, respectively. Tau46 and TauC antibodies were used to stain all the neurons expressing tau protein. We used DAPI (nuclei dye) and tau staining (AT8^+^, Tau46^+^ and TauC^+^) as references to align all antibody staining to the GE section. All staining with the above-mentioned antibodies and DAPI were combined into a single stacking image and assigned into 11 channels in the 10x Genomics Loupe Browser. Then ST spots with AD associated pathology (Aβ^+^, AT8^+^, Aβ^+^/AT8^+^, Aβ^+^/AT8^+^/GFAP^+^, and Aβ^+^/AT8^+^/P2RY12^+^) were chosen to identify gene signatures associated with these pathological changes in AD samples (Fig. 1b).

**Fig. 1.**
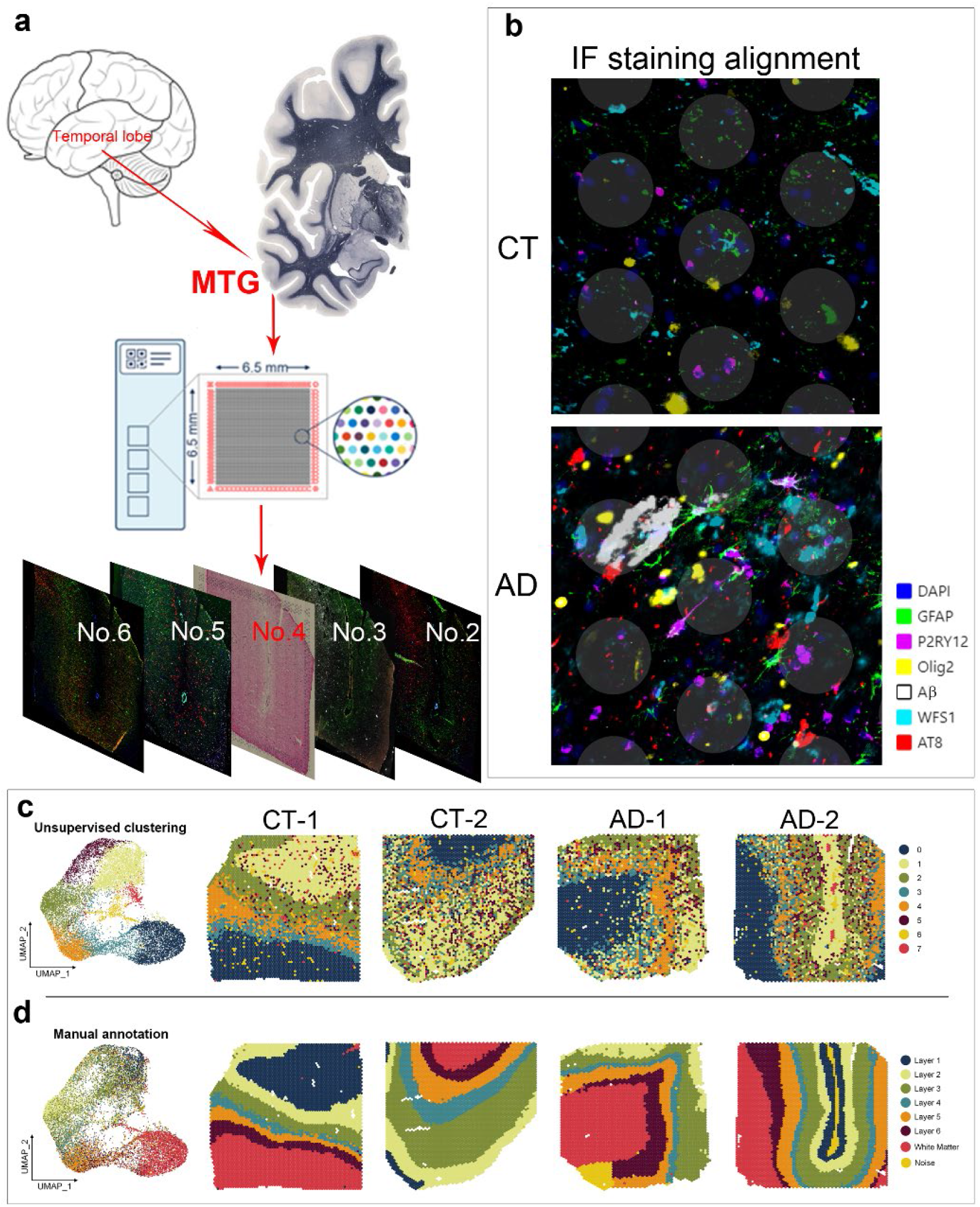
Spatial transcriptomics (ST) of the human middle temporal gyrus (MTG) (**a**) Sequential 10 µm sections of control (CT) and Alzheimer (AD) MTG brain regions were used for all experiments. The middle section (No.4) was used for ST on the Visium platform, and the four adjacent sections (No.2, 3, 5, 6) were used for immunofluorescence (IF) staining. (**b**) Alignment of ST with H&E staining, IF staining of nuclei (DAPI, blue) and cell-type specific markers (GFAP (green), P2RY12 (purple), Olig2 (yellow), WFS1 (teal)) and AD pathological hallmarks (Aβ plaques (white) and AT8^+^ pathological tau (red)). (**c**) Uniform manifold approximation and projection (UMAP) plots show eight clusters (0-7) were identified by the Seurat integration framework using ST spots from both control (CT-1, CT-2) and AD (AD-1, AD-2) human MTG. Spatial maps of the eight clusters for each individual sample from unsupervised clustering. (**d**) Manual annotation of six cortical layers and the adjacent white matter. The left panel figure shows the manually labeled spots (six cortical layers and the white matter) on UMAP space based on the Seurat integration framework. The right panel of the spatial maps show the localization of the manually labeled spots for each individual sample.

### Gene expression in human MTG across cortical laminae and the adjacent white matter

The laminar architecture of the human cortex is closely related to its function and vulnerability to different pathologies^18, 19^. For example, neurons in layers II-III of the temporal lobe including the MTG and the entorhinal cortex are particularly vulnerable to tau pathology in human early AD^6, 7, 20^. Therefore, to identify gene signatures specific to each layer and how they may be differentially expressed in disease, we first assign ST spots to their corresponding layers in each sample, and generated an unsupervised UMAP and the spatial map of each sample with unsupervised clusters (Fig. 1c). Although UMAP can assign a unique cluster to the white matter, it cannot segregate 6 layers in the gray matter of the cortex into individual clusters in the spatial map. In contrast, supervised approach guided by manual annotation of layers based on H&E staining and IF staining of layer-specific markers (see details in the Methods) can clearly defined six cortical layers, unless missing, and the white matter (Fig. 1d).

Based on manual annotation of each layer, ST spots were assigned to their corresponding layer for each sample. Layer-specific ST spots from all four samples were combined as a cluster, and underwent DEG analysis against ST spots in other layers, excluding noise spots. DEGs that were significantly upregulated in individual layers were chosen to quantify their gene expression level across all layers (Supplementary Table 3). In order to identify unique layer-specific marker genes across AD and control brain samples, the DEGs were filtered out if they were not layer-unique DEGs or showed a weak consistency among four samples (consistency level as 2; details can be found in Methods). The layer-specific marker genes (Fig. 2a) were then visualized on spatial maps in Loupe Browser (Fig. 2b). Furthermore, the layer-specific expression of several marker genes were further validated by their in situ hybridization (ISH) data from the Allen Brain Institute’s Human Brain Atlas (Fig. 2b). To verify if the identified layer-specific marker genes can be used to define the laminar architecture of human frontal cortex in publicly available datasets, we probed the expression of our layer-specific marker genes on Visium ST data generated by 10x Genomics (Fig. 2c) and another group^17^ (Supplementary Fig. 1). Overall, we demonstrate that the layer-specific marker genes we identified are powerful in defining the anatomical architecture of the human MTG and the frontal cortex.

**Fig. 2.**
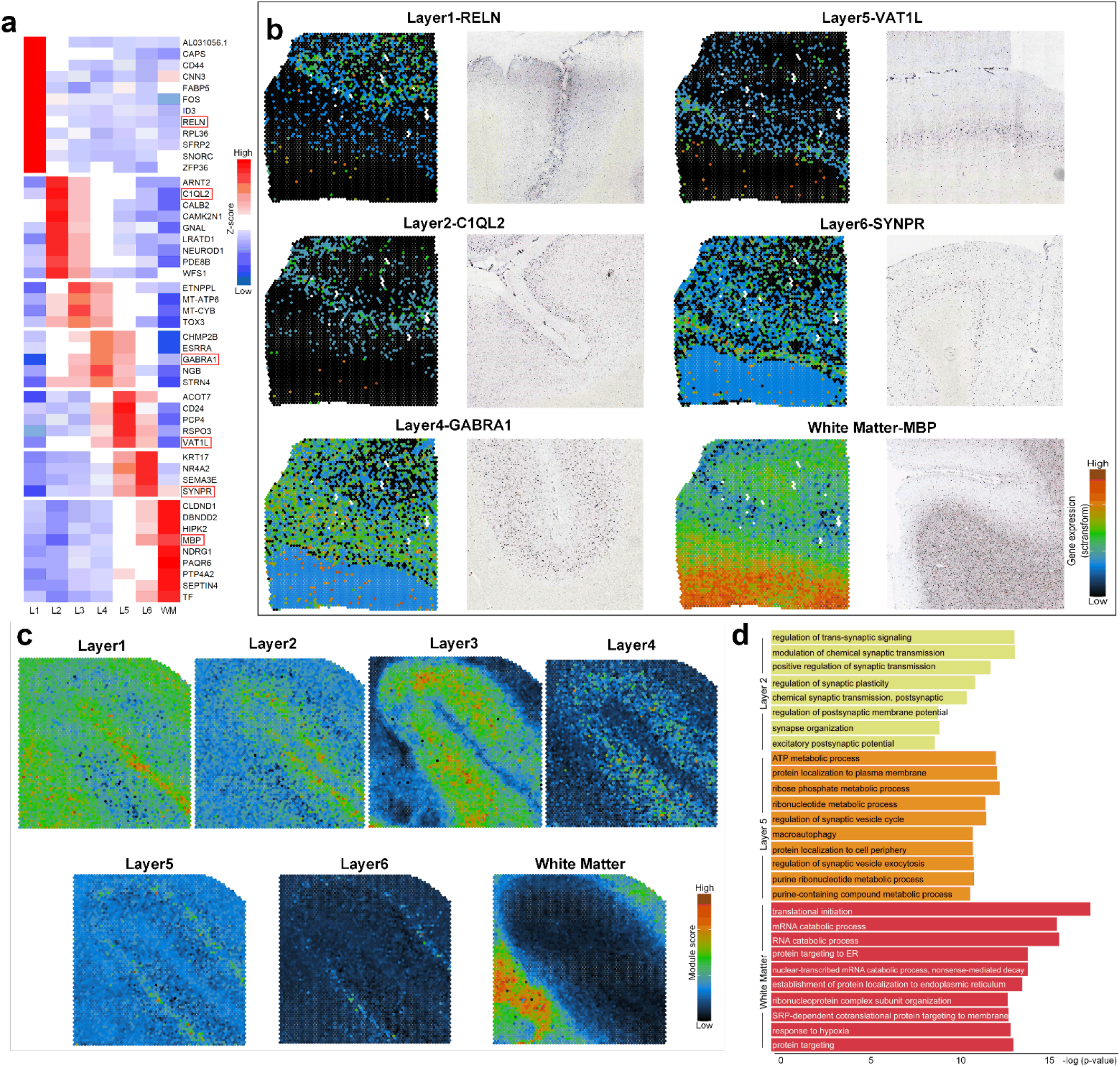
Layer-specific marker genes define the anatomical architecture of the human MTG and the frontal cortex. (**a**) Heatmap of Z-scores for layer-specific marker genes identified from control and AD human MTG. Identified layer-specific marker genes, *RELN* (layer 1), *C1QL2* and *WFS1* (layer 2), *PCP4* and *VAT1L* (layer 5), GABRA1 (layer 4), *SYNPR* (layer 6), and *MBP* (white matter), are boxed in red and were used for spatial map registration using Loupe Browser shown in (**b**). (**b**) Spatial maps of layer-specific marker genes (left panel) in our samples were validated by in situ hybridization (ISH) data (right panel) from the Allen Brain Institute’s Human Brain Atlas (**c**) Public available 10x Visium ST data of the human frontal cortex is annotated by our layer-specific marker genes. (**d**) Gene Ontology (GO) enrichment analysis of upregulated genes in layer II, layer V and the adjacent white matter.

Next, we used the Gene Ontology (GO) enrichment analysis of the upregulated DEGs to define layer-specific biological processes. We found that layers II & III are enriched with synaptic transmission and trans-synaptic signaling, layer V with ribonucleotide metabolic process, regulation of synaptic vesicle cycle, protein targeting and macroautophagy, while the white matter is enriched with RNA catabolic process and protein targeting to membrane and ER (Fig. 2d). Given the important role of synaptic function in the pathogenesis of AD^21, 22^, its enrichment in layers II and III may contribute to the vulnerability of these layers to AD pathology^8, 18, 23, 24^.

In order to investigate the cell type distribution in each cortical layer and the white matter, we performed single cell integration^25^ for our four samples with publicly available snRNA-Seq datasets of human entorhinal cortex, a region in close proximity to the MTG. The results show that excitatory (EXC) neurons are predicated to be mainly located at layers II-V (Fig. 3), which is consistent with the major distribution of Aβ plaques and NFTs in such layers^6, 7, 26, 27^.

**Fig. 3.**
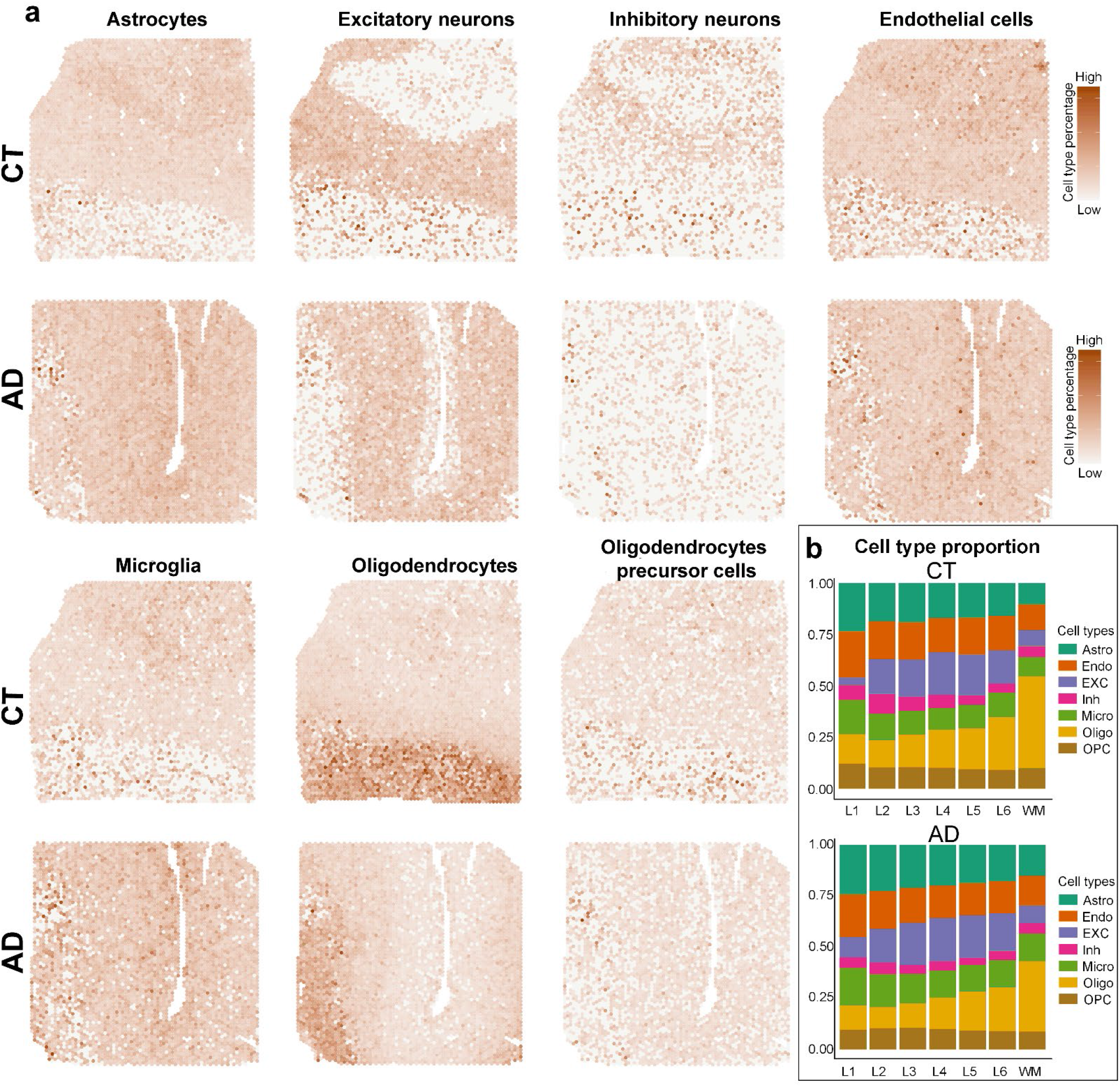
Single-cell integration of snRNA-seq data and Visium ST data from human MTG. (**a**) Spatial maps of the cell-type-specific distribution for astrocytes (Astro), excitatory neurons (EXC), inhibitory neurons (Inh), endothelial (Endo), microglia (Micro), oligodendrocytes (Oligo), and oligodendrocyte precursor cells (OPC) in control (CT) and AD human MTG. Areas enriched for each cell-type are shown in brown. (**b**) Stack bar plots quantifying the average percentage of seven cell types in layers I-VI (L1-L6) and the adjacent white matter (WM).

### DEG analysis between AD and control cases

To better understand how early AD pathology affects the global gene expression in the human MTG, we performed DEG analysis by combining the ST spots in two AD vs. two control samples. We compared the gene expression between AD and control samples, and found 2,164 genes were significantly upregulated while 89 were downregulated in AD samples (Supplementary Table 4). Interestingly, some DEGs have also been identified in recently published bulk RNA-Seq^28^, snRNA-Seq^9, 10, 12^, single-soma RNA-Seq^29^, single microglia RNA-Seq^30^ and the proteomic datasets^31^ from human AD and control cases (Figure 4a). Furthermore, the GO enrichment analysis of these upregulated DEGs reveals many known AD-related biological processes including protein targeting to membrane and ER, RNA catabolic process, viral gene expression, synaptic transmission and trans-synaptic signaling, protein folding and gliogenesis (Fig. 4b). In addition, the GO analysis of the 89 downregulated DEGs reveals exclusive biological processes related to metal ion homeostasis (Fig. 4b).

**Fig. 4.**
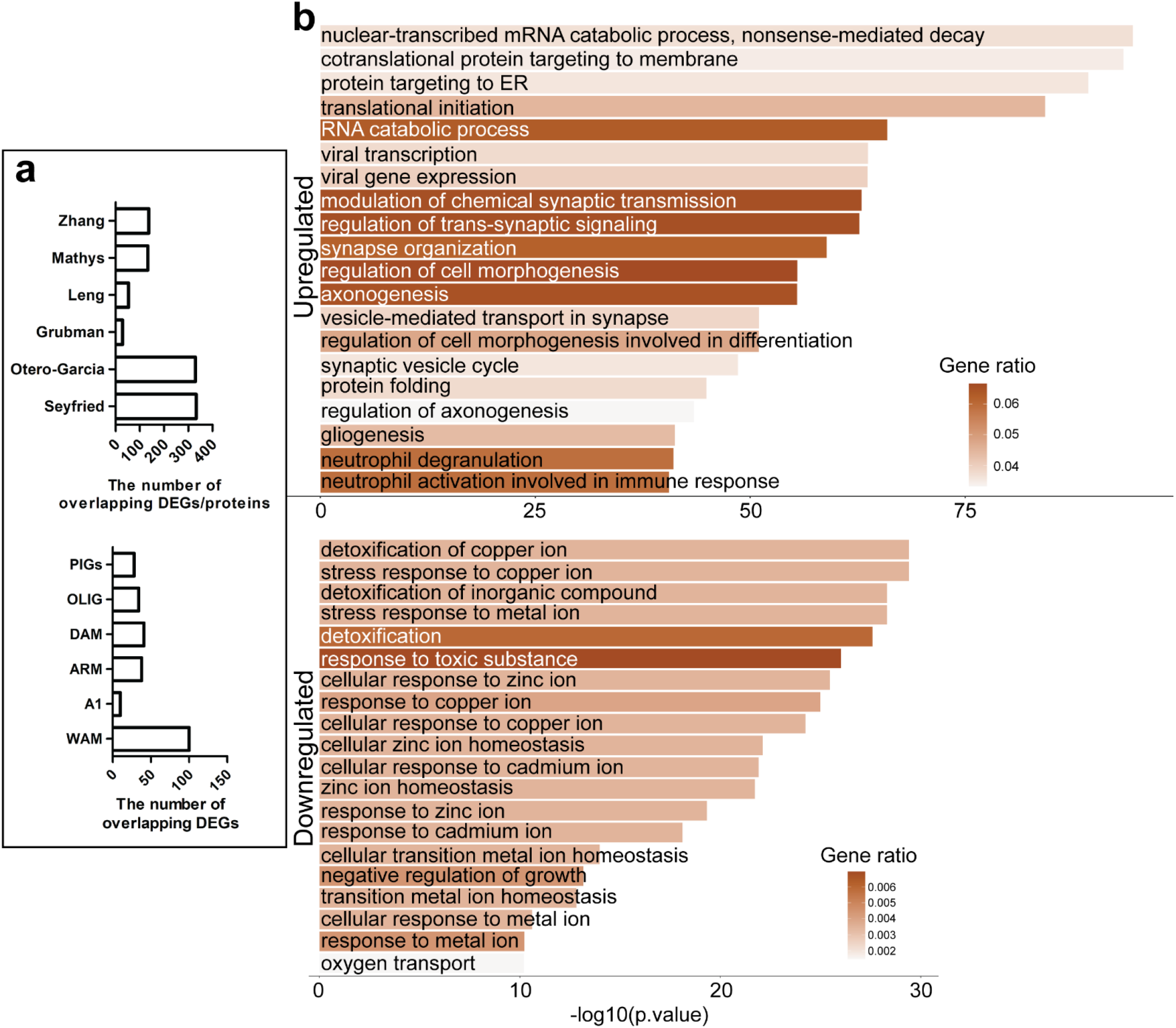
Differentially expressed genes in AD vs Control human MTG. **(a)** Top panel: Overlap of DEGs identified in our study with previous transcriptomic and proteomics datasets published by Zhang^28^, Mathys^10^, Leng^9^, Grubman^12^, Otero-Garcia^29^, and Seyfried^31^. Bottom panel: Overlap of DEGs identified in our study with previously defined AD associated gene modules. **(b)** Gene Ontology (GO) pathway analysis of differentially expressed genes (DEGs) between AD and controls. Pathways for upregulated genes in AD vs control (top) and downregulated genes (bottom) are shown as bar plots on the y-axis, where the x-axis is the negative log-transform of the adjusted p-value. The gene ratio, indicating the number of positive hits, and therefore prevalence of a gene in each pathway is shown in shades of brown, where a darker shade represent a higher ratio. PIG, plaque-induced gene; OLIG, oligodendrocyte gene; DAM, disease-associated microglia; ARM, amyloid-response microglia; A1, A1 astrocytes; WAM, white matter-associated microglia.

### Identification of co-expression gene modules associated with AD pathology

In order to identify gene modules associated with AD pathology, we used weighted gene co-expression network analysis (WGCNA)^16^ to generate modules that best represent similar expression patterns, if any, from our DEGs. We investigated 2,164 upregulated and 89 downregulated genes in AD compared to controls across the full library of 17,203 ST transcriptome profiles and identified four gene modules (M1-4) (Fig. 5, Supplementary Table 5). Based on the RNA-Seq of cell types isolated from human brain^32^, we found that M1 genes are mainly enriched in microglia and astrocytes, M2 in oligodendrocytes, M3 in astrocytes, and M4 in neurons (Supplementary Table 5). Furthermore, the GO enrichment analysis of the component genes in each module reveals that M1 is enriched with biological processes like immune response, synapse pruning and response to unfolded protein; M2 with ensheathment of neurons, myelination and oligodendrocyte differentiation; M3 with lipoprotein particle remodeling, regulation of cell morphogenesis and protein localization to ER; and M4 with synaptic transmission, trans-synaptic signaling and synaptic vesicle cycle (Figs. 5a-d). The enriched biological processes in each module are consistent with its cell-type enrichment. Importantly, 6 of M1 genes and 21 of M2 genes are overlapped with the recently identified PIG module genes and the OLIG response genes^13^, respectively (Supplementary Table 5).

**Fig. 5.**
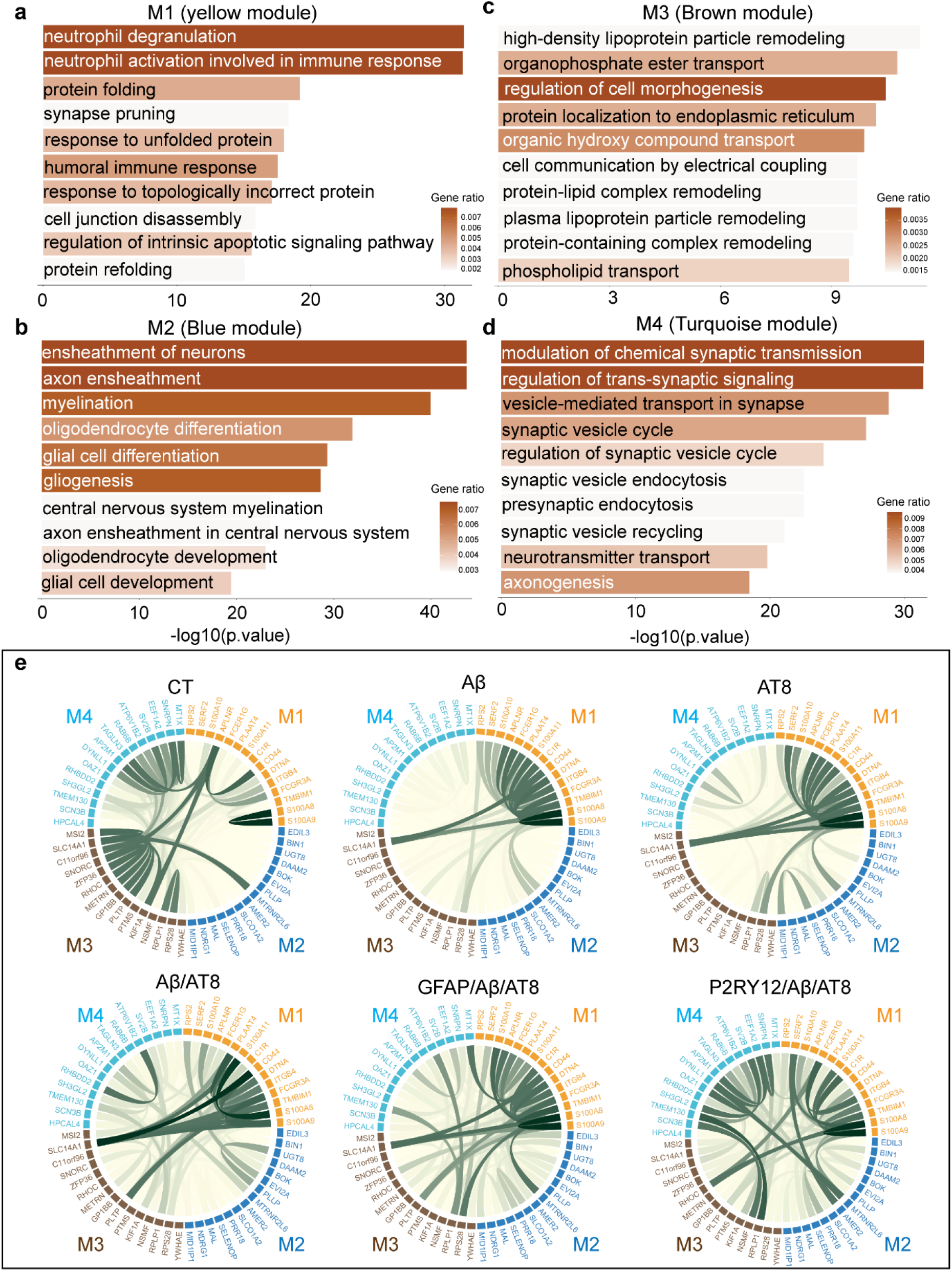
AD-associated gene modules and their co-expression patterns within areas of AD pathology. (**a**-**d**) Gene-ontology (GO) pathway analysis of modules (M1-M4) identified by weighted gene co-expression network analysis (WGCNA) of differentially expressed genes (DEGs) in AD vs control human MTG. Bar plots indicate pathways and functions associated with each module (M1-4) on the y-axis and the negative log transform of the p-value on the x-axis. The gene ratio, indicating the number of positive hits, and therefore prevalence of a gene in each pathway is shown in shades of brown, where a darker shade represent a higher ratio. (**e**) Circos plots of the connectivity strength within and across modules for ST spots in control (CT) human MTG, and with specific AD pathologies (Aβ, AT8, Aβ/AT8, GFAP/Aβ/AT8, and P2RY12/Aβ/AT8). Circos plots are composed of four different modules M1-4 (top 15 hub genes each module); nodes which represent highly deregulated genes from each module; and edges (green lines) which indicate degrees of co-expression (high: dark green, low: light green).

In order to investigate if and how various AD pathologies affect the co-expression of these four gene modules, we performed WGCNA of Top 15 hub genes with the highest closeness centrality score selected from each module (60 genes in total) using only ST spots colocalized with positively stained with either one of the 5 different AD pathologies (Aβ^+^, AT8^+^, Aβ^+^/AT8^+^, Aβ^+^/AT8^+^/GFAP^+^, or Aβ^+^/AT8^+^/ P2RY12^+^) from AD samples. All ST spots from the control non-AD samples were served as a control. Our WGCNA analysis yields connectivity matrixes demonstrating the correlation patterns among those module-specific genes in response to different AD pathologies. The results, visualized by the circos plots showed that the co-expression gene relations within M3 and M4 as well as between M1-3 in controls decrease in the presence of most AD pathologies, whereas the co-expression gene relations within M1 and M2 markedly increase (Fig. 5e). The exception is that the co-expression gene relations within M4 do not decrease in spots with Aβ^+^/AT8^+^/ P2RY12^+^ compared to control spots (Fig. 5e). Also, the co-expression gene relations between M1-4 changed in AD samples compared to control samples (Fig. 5e). Furthermore, the co-expression gene relations of M1-4 change as the distance (45-245 µm) from ST spots with AD pathology increases (level 1: 45 µm; level 2: 145 µm; level 3: 245 µm) (Supplementary Fig. 2).

### Identification of gene signatures and pathways associated with AD pathology

To further identify unique gene signatures and pathways associated with AD pathology (Aβ^+^, AT8^+^, Aβ^+^/AT8^+^, Aβ^+^/AT8^+^/GFAP^+^, or Aβ^+^/AT8^+^/P2RY12^+^), we performed DEG analysis between ST spots with each AD pathology and surrounding level 3 spots (245 µm from the site of pathology). Compared to levels 1 & 2, level 3 spots are of a greater distance from the site of pathology and should exhibit a greater difference in gene expression. It was found that there are many known and novel gene signatures associated with each AD pathology (Supplementary Table 6). The Top-10 upregulated and downregulated genes exclusively associated with each AD pathology were shown as heatmaps (Fig. 6a). We also found that common DEGs and biological processes are shared among a range of AD pathology (Supplementary Table 7). In addition, we found that the comparison between spots with AD pathology and the level 2 spots showed both shared and distinct DEGs and GO pathways with level 3 spots (Supplementary Fig. 3).

**Fig. 6.**
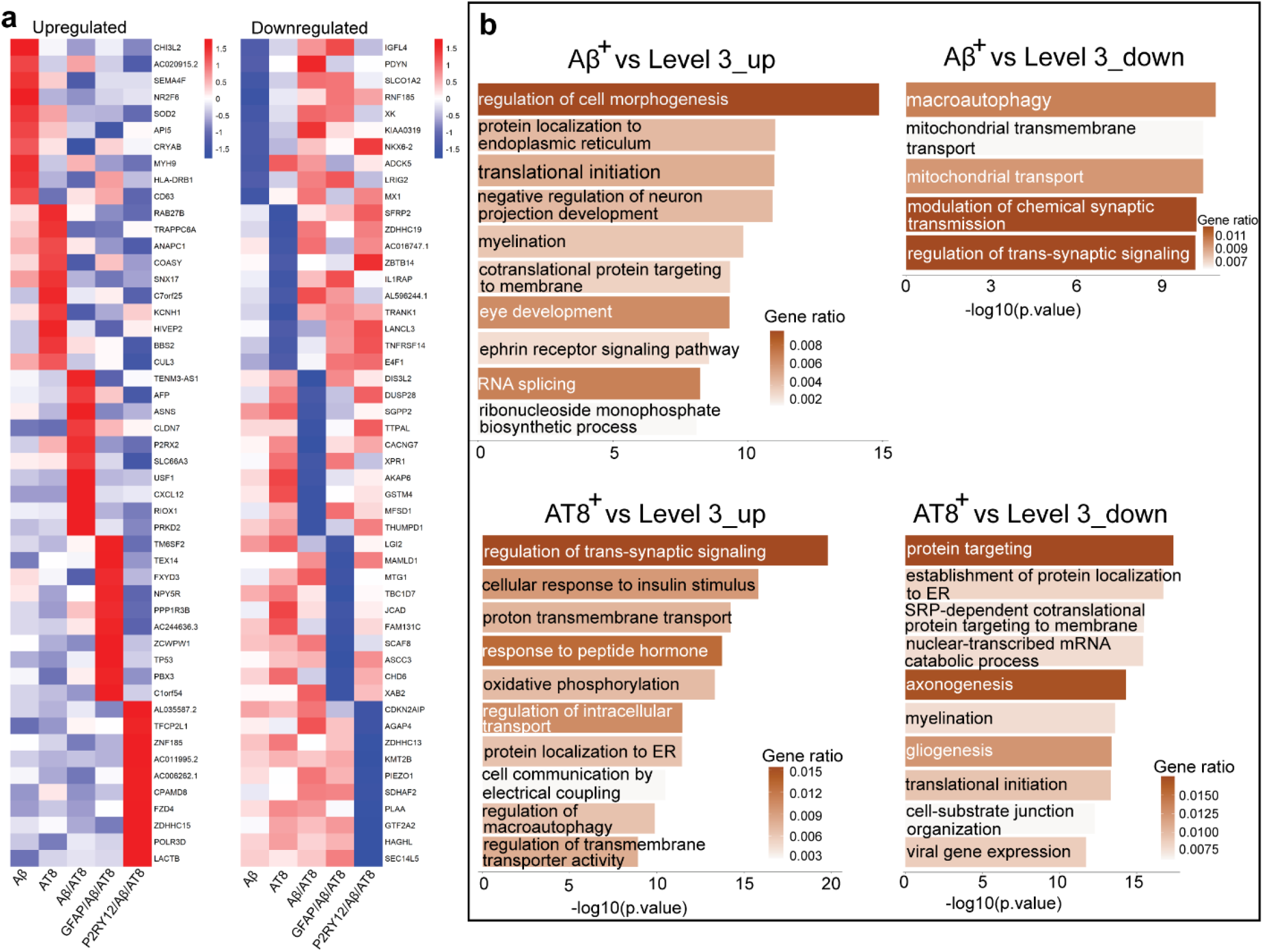
AD associated gene signatures and pathways. (**a**) Heatmap of Z-scores for upregulated and downregulated differentially expressed genes (DEGs) specific to ST spots localized with and without different AD pathologies Aβ, AT8, Aβ/AT8, GFAP/Aβ/AT8, and P2RY12/Aβ/AT8. DEGs were identified from five pathological regions vs. surrounding level 3 spots of two AD samples at the pseudo-bulk level. (**b**) Gene ontology (GO) pathway analysis of identified DEGs specific to ST localized with Aβ plaques (top row) or pathological tau, AT8 (bottom row). Pathways and functions associated with both upregulated genes (left) and downregulated genes (right) are shown.

Next, we determined the biological processes that are enriched in each AD pathology via the GO enrichment analysis. The upregulated pathways associated with Aβ^+^-spots include protein targeting to membrane and ER, translational initiation and myelination, all of which are downregulated in AT8^+^-spots (Fig. 6b, Supplementary Table 6). The downregulated pathways associated with Aβ^+^-spots include the regulation of trans-synaptic signaling, macroautophagy and transmembrane transport, which are upregulated in AT8^+^-spots (Figure 6b, Supplementary Table 6). These results suggest that Aβ^+^-plaques and AT8^+^-tau pathology may induce differential alteration of biological processes, at least in the early stages of AD, although Aβ is believed to be the upstream driver of the tau pathology in genetic forms of AD^33^.

In addition, we quantitated the proportion of spots with AD pathology (Aβ^+^, AT8^+^, Aβ^+^/AT8^+^, Aβ^+^/AT8^+^/GFAP^+^, or Aβ^+^/AT8^+^/P2RY12^+^) in cortical layers and the white matter. It was found that the proportion of spots with AD pathology is much higher in layers II, III and V (Supplementary Table 8). This is consistent with the selective vulnerability of these layers in early AD^8, 18, 23^. Furthermore, the GO enrichment analysis showed dominant biological processes enriched in layers II, III and V are related to synaptic functions (Supplementary Table 3).

### Validation of DEGs associated with AD pathology at the single-cell level using smFISH

Since Visium does not generate single-cell level transcriptomics, we cannot infer the cell-type information of identified DEGs. To solve this issue, we chose RNAscope smFISH in combination with post-IF of Aβ plaques, NFTs (AT8^+^), and GFAP to validate DEGs associated with these AD pathologies at the single-cell level using the same cohort of samples as the Visium ST and a new cohort of 3 early AD cases and 3 control cases. We selected 10 RNAscope probes against cell type-specific marker genes and AD pathology-associated DEGs identified in this study (Supplementary Fig. 4). Consistent with a previous report^32^, *C1QB* and *SPP1* are mainly detected in microglia, *MBP*, *CRYAB*, *CD9* in oligodendrocytes, *GFAP* and *SLC1A2* in astrocytes, NeuN (*RBFOX3*) and *YWHAH* in neurons, and CD63 were detected in both neurons and glia (Figs. 7a-d). Furthermore, we found significantly increased mRNA levels of *C1QB*, *SPP1*, *MBP*, *CRYAB*, *CD9*, *GFAP*, *YWHAH* and CD63, and decreased mRNA levels of *SLC1A2* in AD cases compared to their levels in controls, particularly in those cells within regions with AD pathology (Fig. 7e).

**Fig. 7.**
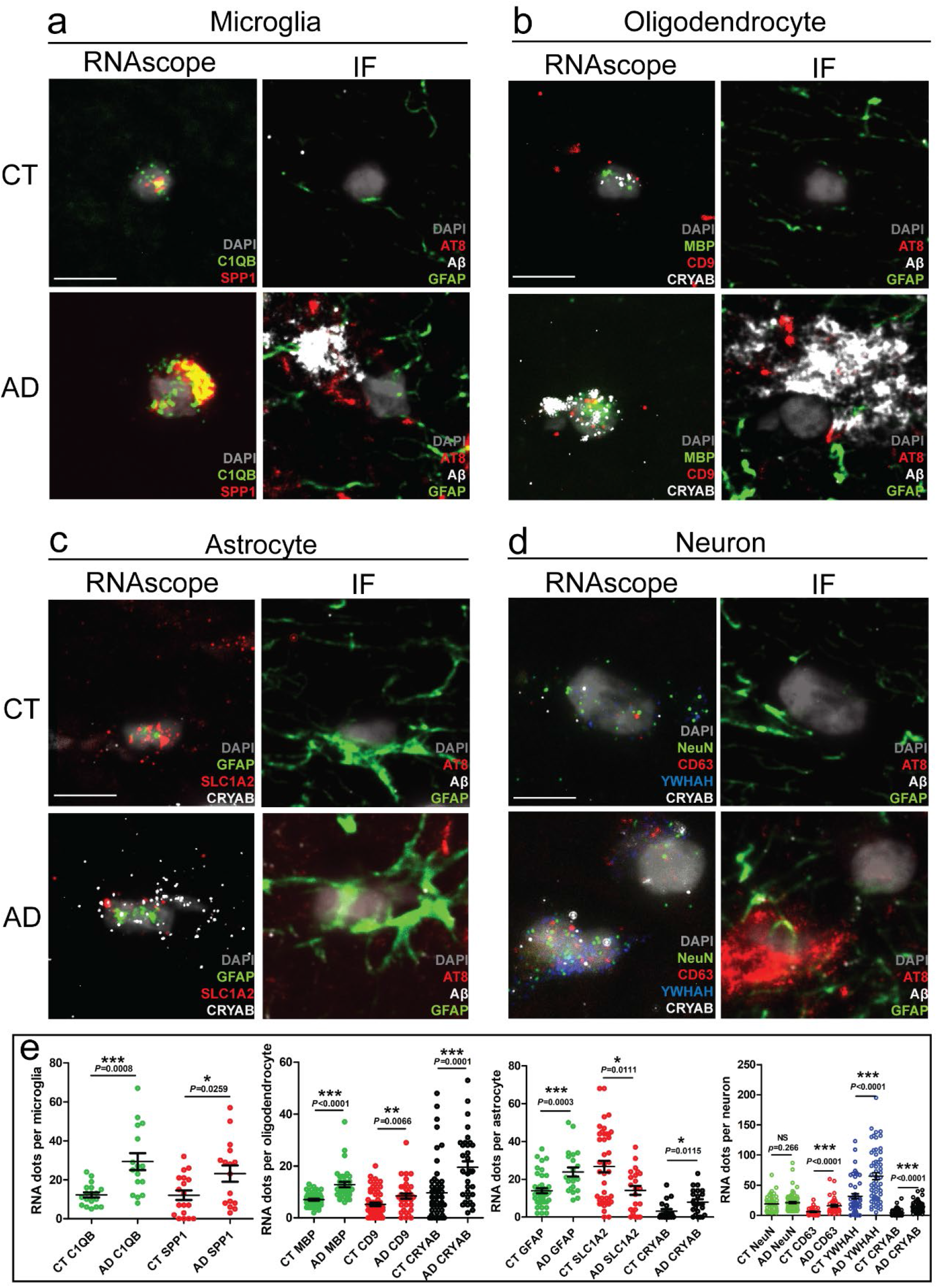
Validation of DEGs associated with AD pathology at the single-cell level using RNAscope smFISH. (**a**-**d**) Representative RNAscope images (left) of 4 different cell types were registered with their corresponding post-Immunofluorescence (IF) Staining (right) in control (CT, top) and Alzheimer (AD, bottom) cases. (**e**) RNAscope probes against human *C1QB* and *SPP1* in microglia (**a**); *MBP*, *CD9* and *CRYAB* in oligodendrocyte (**b**); *GFAP*, *SLC1A2*, and *CRYAB* in astrocyte (**c**); and NeuN (*RBFOX3*), *CD63*, *YWHAH*, and *CRYAB* in neuron (**d**) were quantified and compared between CT and AD. Images of 4 rounds RNAscope and IF were performed on 4 different CT and 4 different AD cases. All images from a same cell were aligned and registered by imageJ. For quantification, regions of interest (ROI) in AD were cropped from regions with abundant Aβ or AT8 staining, while ROI in CT were randomly pricked from different layers. 10 ROIs from each case were collected and all the cells with in ROI were labeled with microglia, oligodendrocyte, astrocyte, or neuron according to the expressions of *C1QB*, *MBP*, *GFAP*, and NeuN. Cells with no labels or multiple labels were dropped during quantification. Overall, we got 5∼10 microglia and astrocytes, and 10∼15 oligodendrocytes and neurons from each case. Except NeuN, all the other probes changed significantly in corresponding cell type as shown. * P< 0.05, ** P<0.01, *** P<0.001 (Mann-Whitney test, CT vs AD).

## Discussion

Our study is among the first to implement Visium ST technology in human AD brain tissue. We define the molecular signatures of cortical laminae and the white matter (Fig. 1d) and identify unique gene signatures (Fig. 2a) and biological process pathways (Fig. 2d) associated with a range of AD pathology by analyzing DEGs within the intact spatial organization of the human MTG from early AD cases and controls. Furthermore, representative genes associated with two main AD pathological markers (Aβ plaques and NFTs) are confirmed at the single-cell level by the method of smFISH, including *CD9*, *C1QB*, *SPP1*, *CD63*, *CRYAB*, *YWHAH*, *SLC1A2*, *MBP*, and *GFAP* (Fig. 7). To be noted, we did not include late AD cases at Braak stages V-VI in this study because we wanted to exclude the downstream gene changes that occur in late AD. We sought to evaluate the transcriptomic changes in the early phase of the disease because we are interested in defining biomarkers of early disease. Previous studies also indicate that major transcriptional changes appear early in AD^9, 10^.

### Identification of novel layer-enriched marker genes in human MTG and reproducibility of these identified marker genes in human cortex with similar architecture

Cortical laminar-specific marker genes in human MTG have been previously identified by in situ hybridization of targeted ∼1,000 genes important for neural functions at a cellular resolution^34^, and by unbiased snRNA-seq at the single-cell level^35^. Recently unbiased ST in the adult human dorsolateral prefrontal cortex (DLPFC)^17^ not only validated laminar enrichment of some canonical layer-specific genes that were previously identified in the rodent^36^ and human cortex^34, 35^, but also identified a number of previously underappreciated layer-enriched genes in human DLPFC^17^. Several genes identified as cell-type markers in specific cortical layers of human MTG are also enriched in the same cortical layers of human DLPFC^17, 34^, suggesting laminar markers are conserved in human cortex with similar layer architecture.

By applying ST in human MTG from both AD and control cases, we identified DEGs and gene clusters that correspond to anatomical layers of human MTG (Fig. 2a). Some cortical layer- and white matter-specific marker genes were consistent with the previous reports^17, 34, 35^, including *RELN* (layer 1), *C1QL2* and *WFS1* (layer 2), *PCP4* and *VAT1L* (layer 5), *SYNPR* (layer 6), and *MBP* (white matter). Importantly, we also identified novel marker genes for layers I-VI and the white matter (Fig. 2a), which were validated by ISH data from the Allen Institute for Brain Science (Fig. 2b) and by annotating the specific cortical layers and the white matter in publicly available Visium ST data of the human frontal cortex (Fig. 2c). This again demonstrates the conservation of laminar markers in human MTG and frontal cortex. The identification of new layer-enriched marker genes will not only help us understand the anatomical architecture of the human cortex but also determine the layer-specific cellular vulnerability to AD pathology (e.g. the excitatory neuronal vulnerability in layers II-III of human temporal cortex^8, 9^ and frontal cortex^17^. Especially since the unique marker genes identified in this study can be used for annotating the cortical layers and the white matter in both healthy controls and AD samples, avoiding potentially confounding effects of AD pathology on gene expression and their corresponding layer in AD samples. Indeed, with the aid of efficient annotation, we found that the number of ST spots with AD pathology is much higher in layers II, III & V compared to other layers (Supplementary Table 8). Also, these two layers are enriched with genes associated with synaptic functions in both AD and control samples (Supplementary Table 3), suggesting that the cortical superficial layers may be vulnerable to AD pathology due to both intrinsic (cell functions) and extrinsic factors (plaques and tangles).

### Comparing the gene signatures, pathways and gene modules identified in this study with those identified by other techniques

Comparing ST spots in two AD samples to spots in two controls, we identified 2,253 (i.e., 2,164 upregulated and 89 downregulated) DEGs (Supplementary Table 4). The GO enrichment analysis of the 2,253 DEGs revealed biological process pathways (Supplementary Table 4) that may contribute to the development and progression of AD pathology. To be noted, Top-10 enriched pathways (Fig. 4b), especially the regulation of trans-synaptic signaling, have also been identified in previous AD datasets^9, 10, 12–14^. Interestingly, most of the downregulated genes are related to metal ion homeostasis, which has been implicated in the pathogenesis of AD^37, 38^.

Furthermore, WGCNA of the 2,253 DEGs identified four gene modules (M1-4), which are surprisingly enriched with gene signatures and pathways mainly associated with microglia, astrocytes, oligodendrocytes and neurons (Figs. 5a-d). Importantly, these modules are also overlapped with several modules identified in previous AD datasets (Fig. 4a). These results suggest that the global gene expression alterations in human AD identified by ST are consistent with previous findings using the bulk RNA-Seq, snRNA-Seq, and large-scale proteomic techniques, which strengthens the feasibility and reliability of implementing ST in studying the molecular mechanisms underlying the pathogenesis of AD.

### The interactions between neurons and glia are altered in the presence of a range of AD pathology

Neurons and glia (microglia, astrocytes and oligodendrocytes) interactions have been found to play important roles in the development and progression of AD pathology, especially Aβ plaques and NFTs^2, 10, 13^. The four gene modules (M1-4) identified by WGCNA in this study correspond to four main cell types in the human brain. We compared the co-expression patterns of these modules in ST spots with AD pathology (Aβ^+^, AT8^+^, Aβ^+^/AT8^+^, GFAP^+^/Aβ^+^/AT8^+^, or P2RY12^+^/Aβ^+^/AT8^+^) in AD cases *vs* all spots in control samples or *vs* adjacent ST spots without corresponding AD pathology in AD cases. We found that the co-expression of microglia- and astrocytes-related M1 genes and oligodendrocyte-related M2 genes were enhanced in the presence of all five AD pathologies, whereas the co-expression of astrocyte-related M3 genes were reduced. The co-expression of neuron-related M4 genes were reduced in most AD pathology except for P2RY12^+^/Aβ^+^/AT8^+^ ST spots (Fig. 5e). We may not have observed obvious changes of M4 in those ST spots due to the fact that P2RY12^+^ microglia are homeostatic and protective microglia^39, 40^, therefore neurons within those spots may not be undergoing degeneration in AD cases with mild-moderate tau pathology (Braak stages III-IV). We identified several hub genes (APLNR, KIF1A, SNORC, TAGLN3, SLC14A1, and MTRNR2L6) of M1-3 in controls, whose strong co-expression disappeared in ST spots with AD pathology in AD cases (Fig. 5e). Interestingly, APLNR and SLC14A1 have also been found to be upregulated in AD^41–44^. The function of KIF1A has been shown to be impaired in the presence of Aβ oligomers^45^. While the repression of TAGLN3 has been recently attributed to APOE4 driven inflammation^46^ and MTRNR2L6 peptides have been identified as neuroprotective species against Aβ toxicity in neurons and other cell types^47, 48^. MTRNR2L6 gene is also found to be one of the most downregulated DEGs in AD patients^49^.All of which indicate that a loss in co-expression of these hub genes may be sufficient to induce differential alterations in biological processes critical for the prevention of neurodegeneration and neuroprotection.

To reduce technical variability and genetic background between AD and controls, we compared the co-expression patterns of these four gene modules in spots with AD pathology *vs* adjacent spots without AD pathology only in AD cases. We detected that the co-expression of M1 genes were gradually reduced, whereas the co-expression of M2 genes are enhanced as the distance from ST spots with AD pathology increases (45-245 µm) (Supplementary Fig. 2). These results demonstrate that the co-expression patterns of neuron- and glia-related gene modules are altered by a range of AD pathology, supporting the hypothesis that interactions between neurons and glia are involved in the development and progress of AD pathology. It should also be noted that not only Aβ^+^ plaques, as previously suggested^2, 13^, but also AT8^+^-NFTs induce changes in neurons and glia co-expression, suggesting that altered co-expression patterns of both neuron-expressed genes and glia-expressed genes are implicated in the pathogenesis of AD.

Differential gene expression analysis between ST spots with AD pathology (Aβ^+^, AT8^+^, Aβ^+^/AT8^+^, GFAP^+^/Aβ^+^/AT8^+^, or P2RY12^+^/Aβ^+^/AT8^+^) and adjacent ST spots (levels 2 and 3) without AD pathology in AD cases, identified upregulated and downregulated genes and pathways associated with each AD pathology. Interestingly, ST spots with Aβ plaques shared a significant upregulation of genes and pathways associated with protein targeting, immune responses, myelination, and a marked downregulation of genes and pathways associated with synaptic plasticity (Fig. 6, Supplementary Fig. 3, and Supplementary Table 6). This finding is consistent with recent transcriptomic, epigenomic, and proteomic analyses^9, 10, 12, 13, 29–31^, highlighting the important roles of gliosis and synaptic dysfunction in AD. Surprisingly, ST spots with AT8^+^-NFTs exhibited opposing alterations in gene expression compared to ST spots with Aβ plaques, i.e., upregulation of genes and pathways associated with synaptic plasticity, and downregulation of genes and pathways associated with protein targeting and immune responses. Whether NFTs cause the degeneration of neurons^50^ or are actually part of a protective response that prevents neuronal death is still unclear^51^. The upregulation of synaptic functions and the lack of enrichment of neuronal cell death and apoptosis pathways in ST spots with AT8^+^-NFTs suggest that neurons with NFTs may exhibit a compensatory upregulation of protective mechanisms against neuronal dysfunction or death in the early stages of AD.

Our results not only validated some of the previously identified Aβ plaques-^13, 14^ and tangle-associated gene signatures and pathways^29^, but also identified new gene signatures and pathways specific for each AD pathology. Therefore, providing new insights for understanding the interactions between neurons and glia in early AD.

### Cell type-specific DEGs associated with AD pathology

Since the diameter of Visium ST spot is 55 µm, which generally includes more than one cell, we further validated DEGs associated with AD pathology at the single-cell level. Using RNAscope smFISH assay, we found that the mRNA levels of *C1QB* and *SPP1* were significantly increased in microglia in AD cases compared to their levels in controls, particularly in microglia within close proximity to regions with AD pathology (Fig. 7). This is consistent with previous reports of increased *C1QB* and *SPP1* in AD using snRNA-Seq^10, 52^ and single-microglia RNA-Seq^53, 54^. The complement system, especially C1q, and microglia are activated in early AD, which have been found to mediate early synapse loss in Alzheimer mouse models via the synaptic pruning^55, 56^. The SPP1 (osteopontin) is found to be one of the most upregulated genes in microglia in AD^10^. Although the function of SPP1 in the pathogenesis of AD is unclear, previous reports suggest that it is a chemoattractant and an adhesion protein involved in wound healing and has been identified as a gene related to amyloid-response microglia (ARM)^53^. SPP1 is also found to be an inflammatory marker upregulated in vascular diseases as well as in AD^57^.

*MBP*, *CRYAB*, and *CD9* genes were also found to be significantly increased in oligodendrocytes in AD cases, with even higher expression in oligodendrocytes within regions with AD pathology (Fig. 7). Interestingly, CRYAB (HSPB5) has been found to mediate the inhibition of amyloid nucleation, fibril binding, and fibril disaggregation^58^. The upregulation of oligodendrocyte-related genes has been demonstrated in AD-like animals and human AD at the early stages of the disease^10, 13, 30^. The findings of oligodendrocyte changes from this study and other groups support the important function of oligodendrocytes in early AD. Importantly, *SPP1*, *C1QB* and *CD9* have also been identified in the gene expression profiles of mouse microglia around Aβ plaques. This profile has been variously labeled as disease-associated microglia (DAM)^39^, microglial neurodegenerative phenotype (MGnD)^40^, ARM^53^ or PIG^13^.

In regards to the astrocyte-associated genes GFAP and SLC1A2, we found the former significantly increased, whereas the latter decreased in AD, especially within the regions with GFAP^+^/Aβ^+^/AT8^+^-staining (Fig. 7). In line with our findings, previous reports have widely demonstrated the upregulation of GFAP and downregulation of SLC1A2 in AD-like animals and human AD compared to controls^9, 59, 60^. The changes of these two genes together with the overlap of our identified DEGs with previously identified genes associated with A1 astrocytes^59^ indicate that the homeostatic function of astrocytes are reduced in AD, resulting in the accumulation of reactive astrocytes (gliosis) and immune responses around AD pathology.

The *YWHAH* mRNA is enriched in neurons and its expression level is significantly increased in AD (Fig. 7). This gene encodes the 14-3-3 protein, which is recently found to be increased in tangle-bearing neurons and may be involved in the neurodegeneration induced by tau aggregation^29^. It should be noted that 14-3-3 proteins (YWHAH, YWHAB, YWHAQ, and YWHAZ) have been found in NFTs^61^, and the latter three of them stimulate the phosphorylation of tau and the formation of NFTs^62^. The role of YWHAH, however, is unknown.

### Limitation of this study

Although 10x Genomics Visium ST platform is a powerful technique of characterizing the spatial transcriptomic profiles of rodent and human brain samples, there are a few limitations of this study. First, the Visium ST platform is not at single-cell resolution^13, 14^, although we validated key representative genes at the single-cell level in this study. Thus, we cannot exactly provide the cell-type information for most DEGs identified in each ST spot. Second, we performed the immunostaining of cell type-specific markers and AD pathological markers on adjacent brain sections, but not on the same section used for the H&E staining and the Visium ST assay. Although the distance between the Visium section and the immunostaining sections is 10 or 20 µm, alignment of those sections may produce minor errors in terms of the exact location of each ST spot in adjacent brain sections. Third, although our results from small size samples identified novel and known gene signatures and pathways that are associated with AD pathology, further ST analyses of larger sample size, sex difference and additional functional studies are warranted for fully understanding the relationship between transcriptional changes and various AD pathology. Fourth, we did not focus on particular forms of Aβ and tau aggregates in this study. Further ST studies in combination with immunofluorescence staining of soluble *vs* insoluble forms of Aβ and tau aggregates will be needed to distinguish the transcriptional changes in the presence of various forms of AD pathology.

Overall, our ST analysis highlights the molecular composition of anatomical architecture of the human MTG and the complexity of glial-neuronal interactions in response to a range of AD-associated neuropathology. Although we cannot distinguish drivers of degeneration from compensatory responses, our spatial transcriptomic data provide a resource for exploring the molecular mechanisms underlying the pathogenesis of AD and a platform for discovering potential biomarkers and disease-modifying therapeutics.

## Online Methods

### Human postmortem brain tissues

Human fresh frozen brain blocks were provided by the Arizona Study of Aging and Neurodegenerative Disorders/Brain and Body Donation Program at Banner Sun Health Research Institute^63^ and the New York Brain Bank at Columbia University Medical Center^64^. The demographics of human cases used in this study are listed in Supplementary Table 1. These specimens were obtained by consent at autopsy and have been de-identified and are IRB exempt so as to protect the identity of each patient. Frozen sections (10 μm) were cut from frozen blocks under RNase-free conditions.

### Sample selection and preparation for ST

Thirty milligram brain tissue was homogenized in 500 µl TRIzol RNA isolation reagent (ThermoFisher, Cat# 15596026). RNA was extracted from each homogenate by following the TRIzol RNA extraction procedure. The RNA quality of each sample was assessed by RNA integrity number (RIN) via Agilent 2200 TapeStation system. Samples with a RIN higher than 6.0 were selected and cut into 10 µm sections, which were subjected to IF staining of total tau (TauC antibody) and phosphorylated tau level (AT8, pS202/T205 tau antibody). Regions with the highest TauC^+^ and/or AT8^+^ staining were scored within 6.5 mm × 6.5 mm selected square area.

The time course of the tissue permeabilization for each sample was determined by 10x Genomics Visium Spatial Tissue Optimization (STO) Kit (Slide Kit, Part# 1000191. Reagent Kit, Part# 1000192). Briefly, sections from selected square areas of the brain tissue were mounted on the capture areas of STO slides. Sections were fixed in the -20°C prechilled methanol for 30 min and subjected to H&E staining. The 20x tiles image for each sample was obtained by Zeiss Axio Observer microscope. After imaging, sections were permeabilized with permeabilization enzyme for varying amounts of time, and the reverse transcription (RT) was performed directly on the slide using Cy3-nucleotides. After RT, sections were imaged and aligned with H&E images. The best time course was chosen for ST if its image shows the strongest Cy3 signal and the minimum signal diffusion.

### ST processing

Each optimized human postmortem brain tissue was cryosectioned at 10 µm continuously for seven sections and was labeled sequentially as No.1 to No.7. The middle section (No.4) was mounted on the Visium Gene Expression (GE) slide for ST profiling, sections No. 2, 3, 5, 6 were used for IF staining of cell-type markers and AD pathology. The outmost sections (No.1 and No.7) were used for the RNAscope HiPlex assay. ST were performed using the 10x Genomics Visium Spatial Gene Expression Kit (Slide Kit, Part# 1000188. Reagent Kit, Par# 1000189). The procedures prior to RT are the same as described for STO. For RT, cDNA was synthesized using nucleotides without the label of Cy3. By using Template Switch Oligo, the second strand cDNA (ss cDNA) was synthesized according to cDNA templates captured on the poly T probes. The ss cDNA was then denatured by 0.08 M KOH, washed off, and then amplified using PCR. The cDNA quality control was performed using an Agilent Bioanalyzer high sensitivity chip. After the cDNA concentration was determined, the sequencing library of each sample was constructed using the 10x Library Construction kit (Part# 1000196). Briefly, optimized cDNA was obtained by enzymatic fragmentation and size selection. P5, P7, i7 and i5 sample indexes and TruSeq Read 2 were added via End Repair, A-tailing, Adaptor Ligation, and PCR. The cDNA library with correct sizes was then selected using the SPRIselect reagent (Beckman Coulter, Part# B23318). In order to meet the required sequencing depth, cDNA libraries from four samples were pulled in a NovaSeq6000 SP v1.0 flowcell and paired-end sequencing were performed on an Illumina NovaSeq6000 sequencer at the Genomics Services Laboratory at Nationwide Children’s Hospital.

### IF staining

As described above, sections No.3&5, 2&6, and 1&7 were 0 µm, 10 µm, and 20 µm apart from the GE section, respectively. Section No.3 was sequentially stained with P2RY12/GFAP/AT8, section No.5 with Olig2/Tau46/Aβ, section No.2 with WFS1/AT8/GAD1, and section No.6 with TauC/ENC1/GAD1. Sections No.1&7 were used for the RNAscope HiPlex assay. IF staining was performed as previously described^11^. All sections were air dried at 60°C for 10 min and then fixed and permeabilized by prechilled acetone at -20 °C for 15 min. For sections No.3&5, slides were immersed in 1x PBS for 3 hours at 37°C for antigen retrieval. For sections No.2&6, antigen retrieval was performed by incubating slides in 10 mM sodium citrate (pH6.0, 95°C) for 12 min. Antibodies were incubated sequentially to avoid false-positive results coming from co-immunostaining. On day 1, after 1-h blocking by 10% donkey serum (in 1x PBS), P2RY12 (BioLegend, Cat# 848001. 1:1000) and Olig2 (R&D systems, Cat# AF2418. 1:500) primary antibodies were applied on section No.3 and 5, respectively, overnight at 4°C, while GAD1 (R&D systems, Cat# AF2086. 1:100 in PBS with 10% donkey serum) antibody was applied to sections No.2 and 6 overnight at 4°C. On day 2, after three washes with 1x PBS, secondary antibodies were incubated with corresponding sections for 2 hours at 37°C. The nuclei were stained with Hoechst33342. Autofluorescence was quenched with 0.5x TrueBlack solution in 70% ethanol for 10 min. The coverslips were mounted with Fluoromount-G Mounting Medium (SouthernBiotech, Cat# 0100-01), and the slides were then imaged by Zeiss Axio Observer microscope. Following 1-h re-blocking with 10% donkey serum in 0.3% PBST, primary antibody combinations for each section were incubated at 4°C in the same way as described above for regular IF staining.

### H&E and IF staining image alignment

The 20x tiles images taken by Zeiss Axio Observer microscope were exported into merged and individual channel images by Zen software (v3.2 blue edition). The merged channel images were landmarked according to hematoxylin staining in corresponding H&E images using Fiji “Multi-points” tool. Multi-points landmarks in each merged channel image were then saved as selection and transferred to individual channel images. Image alignment for each channel to H&E was performed using the “Transform/Landmark correspondences” plugin in Fiji. H&E image and transformed channel images were stacked into .tiff image and imported into Loupe Browser (v5.0.1) and Space Ranger (v1.2.2) for further alignment with ST spots.

### Unsupervised and supervised annotation of cortical layers and the white matter of the human MTG

To assign ST spots to their corresponding layers, we first combined all 17,203 spots from our four samples and generated a cross-sample unsupervised Uniform Manifold Approximation and Projection (UMAP) via Seurat (v.3.2.2)^65^ to visualize two dimension-reduced data. All spots were clustered into eight groups and visualized in the original sample using eight colors assigned to each cluster. Although UMAP can roughly reflect the cortical layers’ information and separate the white matter from the gray matter well, it is not powerful to segregate six layers in the gray matter of cortex. Since certain layer(s) may be missing during the brain dissection or mounting brain sections on the GE slide, arbitrarily assigning clusters from unsupervised UMAP into a sample may not be accurate without manual annotations. Moreover, the pathology development in AD brains may also change the gene expression profile and introduce more confounders in clustering. To solve this problem, we used both UMAP and IF staining as references to manually label each cortical layer and the white matter.

First, a UMAP plot for each sample was generated by Loupe Browser to avoid batch effects across samples. In order to assign UMAP spots to corresponding layers, previously identified layer-specific marker genes were chosen to highlight regions of certain layers on the UMAP. A group of marker genes for each layer was shown as a feature combination using a method of Feature Sum. The outstanding spots with higher expression of feature combination genes were selected and temporally labeled with that layer. The boundary of each layer would then be verified in spatial images with different IF staining, which were used as the ground truth. For example, three previously defined layer 2 markers, C1QL2, RASGRF2, and WFS1, were shown on the UMAP^17, 34^. The highlighted spots with higher expression of those three genes were assigned to layer 2 in the category of Layer. Those spots would then be visualized on the image with WFS1 staining to define a clear boundary between layer II and layer I/III. Using the same strategy, the gene expression of RELN and the IF staining of GFAP (astrocytes) were used to define layer 1. Similarly, the gene expression of MAPT, MBP and MOBP (oligodendrocytes), and the IF staining of total tau via TauC and Tau46 as well as the IF staining of Olig2 (oligodendrocytes) were used to define the gray and the white matter, respectively. Since layer IV is an internal granular layer and a distinct layer with dense nuclei that could be identified in both H&E and DAPI staining, we also include the nucleus staining from H&E and DAPI as ground truth when using the canonical layer 4 marker genes (e.g., RORB and PDYN) to label layer IV. In addition, PCP4 was used as a marker gene for layer V, while PCDH17, TLE4 and FOXP2 were used as marker genes for layer VI. Although no IF staining is available in layer V and layer VI, the boundary between those 2 layers are already clear enough by only using layer-specific marker genes. All the spots across four samples were allocated to a certain layer or assigned as noise if their definition was too vague based on UMAP or IF staining. A total of 250 spots were dropped as the noise across four samples.

### FASTQ generation, alignment, and quantification

Raw sequencing data of four human MTG samples were obtained from Visium (BCL files) and processed with the SpaceRanger (v.1.2.2) software to generate FASTQ files via the SpaceRanger *mkfastq* function. FASTQ files were then aligned to and quantified to the expression matrix by using GRCh38 Reference-2020-A and the spaceranger *count* function. The two functions spaceranger *mkfastq* and *count* were used for demultiplexing sample and transcriptome alignment using default parameters.

### Data processing

Seurat (v.3.2.2) was used for the following analysis. For a sample, the spatial file and HDF5 file obtained from spaceranger were loaded into Seurat by the function *Load10X_spatial*. Then we used the recommended sctransform method^66^ to normalize counts by the function *SCTransform* in its default settings. Multiple manual annotations (meta information) were added to each Seurat object via the function *AddMetaData*. All the spots labeled as noise were removed, and the remaining spots were used for the following analyses.

### Data Integration, dimension reduction, spot clustering, and visualization

The Seurat integration framework was used for identifying clusters and reducing confounders from the four samples (two AD samples and two normal tissues). First, 3,000 features were selected for integrating four datasets via setting *nfeature* as 3,000 from the function *SelectIntegrationFeatures.* Then the function *PrepSCTIntegration* and *FindIntegrationAnchors* were used for preparing anchor features and generating anchors for the following analyses. Principal Component Analysis (PCA) was utilized for dimension reduction and denoising via *runPCA*. The top ten principal components (PCs) were select to perform UMAP via runUMAP in the default settings. To better match the six layers and white matter of human cortex structure, we set the resolution as 0.3 to generate spot clustering results by the functions *FindNeighbors* and *FindClusters*. The UMAP plots and spatial maps were generated by *DimPlot* and *SpatialPlot*.

### WGCNA of DEGs between AD and the control

The R package WGCNA (v1.69)^67^ was used to identify gene modules and build unsigned co-expression networks, which include negative and positive correlations. The identified AD upregulated and downregulated DEGs at the pseudo-bulk level was used to identify gene modules. Soft power 3 was chosen by the WGCNA function *pickSoftThreshold*. Then, the function *TOMsimilarityFromExpr* was used to calculate the TOM similarity matrix via setting power = “3”, networkType = "unsigned" and TOMType = "unsigned". The distance matrix was generated by substrating the values from similarity adjacency matrix by one. The function flashClust (v.1.01)^68^ was used to cluster genes based on the distance matrix, and the function *cutreeDynamic* was utilized to identify four gene modules (yellow: M1, blue: M2, brown: M3, and turquoise: M4) by setting deepSplit =3. To determine key genes with the highest degree, the function *TOMsimilartyFromExpr* was used to generate the similarity matrix based on all genes of each module. The matrix was then used for calculating the edges’ closeness centrality score via function *graph_from_adjacency_matrix* and *closeness* from the R package igraph (v.1.2.6) for four modules. To fairly compare closeness scores among different modules, the min-max normalization method was used to normalize closeness score. Based on the normalized closeness score, genes were sorted, and the Top-15 genes were selected to generate four modules circos plot via the circlize package (v.0.4.13)^69^.

### DEG and GO enrichment analysis

The DEG analysis was conducted by the Seurat function *FindMarkers* via grouping AD and control samples (*min.pct* was set as 0.01 and *logfc.threshold* was set as 0.1). The DEGs were selected if the adjusted *p*-value was less than 0.05 and the absolute value of log-fold change was higher than 0.1. Based on the identified DEGs, the enrichment analyses of GO terms (Biological Process), KEGG, and Reactome were performed via the R package clusterProfile (v.3.18.0)^70^ and ReactomePA (v.1.9.4)^71^ using the functions of *enrichGO*, *enrichKEGG*, and *enrichPathway*. The enrichment analysis results were filtered out if the adjusted *p*-value was greater than 0.05. For KEGG analysis, gene database Org.Hs.eg.Db was used for transferring SYMBOL to ENREZID via function *bitr*. R package ggplot2 (v.3.3.2) was used for the visualizations. The sidebar legend was colored by the gene ratio defined as hit genes over the query genes via *enrichGO* function. For each identified gene module, the component genes were used for pathway enrichment analysis.

For each sample, DEG analysis for pre-defined layers was also conducted. Layer-specific DEGs were merged from four samples for layers I-VI and the white matter. We calculated the number of shared DEGs across four samples. Then layer-specific markers were determined by their uniqueness (i.e., unique to one layer in at least two samples compared to other layers’ DEGs) and their consistency (i.e., one DEG being identified from at least 2 samples simultaneously). Enrichment analyses of GO terms (Biological Process), KEGG, and Reactome were performed for DEGs with consistency numbers greater than 2.

### RNAscope HiPlex assay in combination with post-IF staining of Aβ/AT8/GFAP

In order to validate and quantitate the gene expression changes around AD pathology-related regions (Aβ^+^, AT8^+^, and/or GFAP^+^), we performed the smFISH assay using the RNAscope HiPlex12 Ancillary Kit (Cat. No 324120) to simultaneously detect 10 different RNA targets selected from DEGs associated with Aβ plaques and/or AT8^+^-tangles. The process was performed according to the manufacturer’s user manual with a few modifications. Fresh frozen sections from another set of human AD and control brain tissues, as well as sections No.1 & 7 preserved from 10x Visium ST experiment, were fixed in 4% PFA at room temperature for 1 h. The sections were washed twice by 1x PBS and dehydrated sequentially in 50%, 70% and 100% ethanol for 5 min. After permeabilization by protease IV, the sections were incubated with the premixed 10 probes in the ACD HybEZ II oven for 2 h at 40°C, and three amplification steps in 40°C were extended to 45 min instead of 30 min. Since the Alexa 750 channel is not available on our microscope, we changed the original three rounds reaction (four probes per round) to four rounds with only three probes in each round. After each round, the nucleus were counterstained with DAPI for 30 s, and the lipofuscin autofluorescence was quenched by 1 min of 0.5x True Black. The slides were mounted with Fluoromount-G Mounting Medium, and 40x Tiles image were taken by Zeiss Axio Observer microscope after each round. Before the next round, signals were quenched using cleaving buffer and washed twice with 1x PBS containing 0.5% of Tween20. To define the regions with Aβ or AT8, IF staining was performed after RNAscope HiPlex12 assay. Briefly, the signal from round 4 was quenched using the cleaving buffer. After that, the antigen retrieval and staining process were performed as described above. Images from each round and the IF staining were aligned using HiPlex Image Registration Software.

### Spatial integration of snRNA-Seq and ST data from human MTG

The snRNA-Seq datasets from human entorhinal cortex (EC) of AD and controls^9^ were used for performing single-cell registration for our four samples based on R package SPOTlight (v.0.1.6)^25^. The EC datasets were downloaded from https://www.synapse.org/, and the access number was syn22722817. To fairly and precisely integrate the EC datasets and our samples, single-nucleus data labeled as disease stage 0 was selected for the following analysis. The function FindAllMarkers was utilized to identify the EC dataset’s DEGs based on benchmarking labels (Supplementary Table 9). An in-house database named scREAD^72^ was used for acquiring cell-type-specific markers. Final cell-type-specific markers (CTS, Supplementary Table 10) were identified by comparing the overlapped DEGs based on the EC dataset and the markers from scREAD. Next, the proportion of each cell type was estimated by the function spotlight_deconvolution via inputting the defined new CTS markers. The mean proportion of each cell type in each layer and the five pathological regions was estimated and visualized using the R package ggplot2 suite.

### Statistical analysis

No statistical methods were used to predetermine sample sizes, since (*i*) a rigorous statistical framework for design of ST experiments is missing in the literature and (*ii*) existing tools for the design of single-cell genomic experiments are not appropriate for ST experiments and do not consider key factors of ST experiments. In our study, each brain sample has over 4,000 spots and ∼3500 genes per spot, which provides sufficient power for DEG analysis of Visium ST datasets. Prism 5 software was used to analyze the data. All data are expressed as mean ± s.e.m. We performed the D’Agostino–Pearson omnibus normality test to determine whether the data were normally distributed, or the *F* test to determine whether the data assumed equal variances. We then chose the following statistical tests. Nonparametric Mann-Whitney tests were used to compare the number of single RNA dots in each cell human non-AD and AD. All results represent two-sided tests comparing groups of biological replicates. *P* < 0.05 was considered statistically significant for all measures. The *n* values represent the number of spots or cells in each group; exact values are indicated in figure legends.

### Code availability

All the codes used in this study are freely available after being accepted for publication.

### Contributions

H.F. and Qin Ma jointly designed and supervised the study, discussed the results, and wrote the paper. S.C. designed and performed experiments and analyzed the data. Y.C. generated all the pipelines for analyzing the 10x Genomics Visium datasets. L.L., D.A., C.M. and C.W. helped with the data analysis. D.J. and M.E.H. helped with the RNAscope HiPlex assay. G.E.S. and T.G.B. characterized the brain tissues. All authors discussed the results and contributed to the manuscript writing and/or editing.

## Acknowledgements

This work was supported by awards K01-AG056673 and R56-AG066782-01 (H.F.) from the National Institute on Aging of the National Institutes of Health and the award R01-GM131399 (Q.M.) from the National Institute of General Medical Sciences. The work was also supported by the award of AARF-17-505009 (H.F.) from the Alzheimer’s Association, the W81XWH1910309 (H.F.) from the Department of Defense, and the 10x Genomics 2021 Neuroscience Challenge award. We thank Karen Duff from the Columbia University and the University College London, and the 10x Genomics technical support team for helpful discussions. We also thank Amanda Toland, Pearlly Yan, Tom Liu, Jennifer Mele from the Ohio State University and Amy Wetzel from the Nationwide Children’s Hospital for helping with the RNA quality control and the sequencing. Human de-identified brain tissues were kindly provided by the Banner Sun Health Research Institute Brain and Body Donation Program, supported by NIH grants U24-NS072026 and P30-AG19610 (TGB), the Arizona Department of Health Services (contract 211002, Arizona Alzheimer’s Research Center), the Arizona Biomedical Research Commission (contracts 4001, 0011, 05-901 and 1001 to the Arizona Parkinson’s Disease Consortium) and the Michael J. Fox Foundation for Parkinson’s Research and the New York Brain Bank at Columbia University Medical Center. This work used the high-performance computing infrastructure at the Ohio State University.

## Data availability

The raw data and unfiltered UMI count matrices will be available on the Gene Expression Omnibus (GEO), and the processed data can be downloaded at Synapse.org after being accepted for publication. All data and analysis will also be added into our in-house scREAD database at https://bmbls.bmi.osumc.edu/scread/ after being accepted for publication.

---

**Supplementary Fig. 1.**
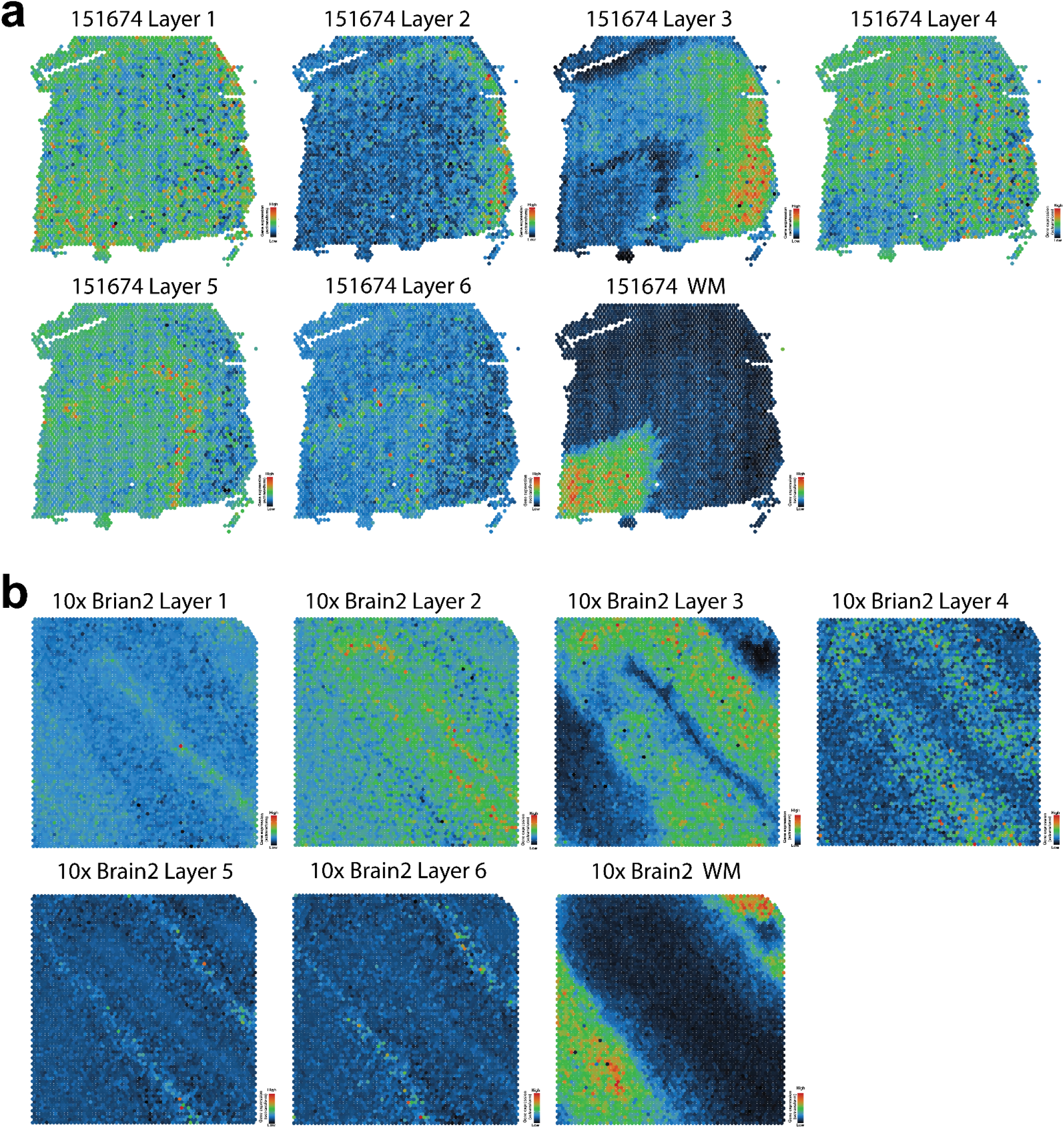
Validation of layer-specific gene modules on public available Visium ST datasets. Spatial maps show the gene module score for each layer-specific marker on the human brain sample from another group^17^ (**a**), and human brain 2 in 10X Genomics’ public dataset (**b**). The red color represents a higher score, indicating the average gene expression value of the gene module higher than random control genes. Conversely, the black color represents a lower score, indicating the average gene expression value of the gene module lower than random control genes. WM, white matter.

**Supplementary Fig. 2.**
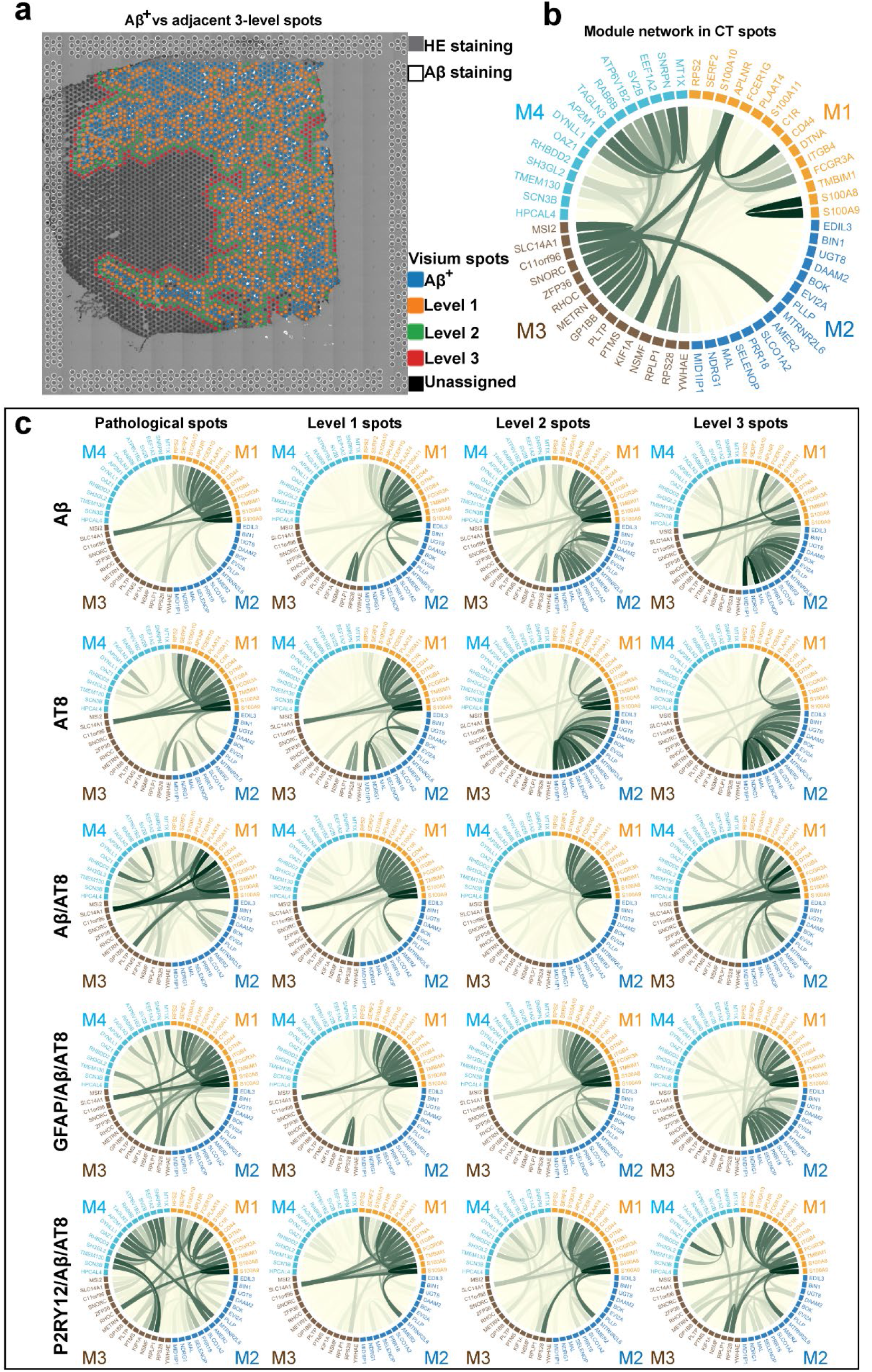
Gene modules associated with AD pathology at varying spatial distances. (**a**) Aβ plaques were stained with human specific Aβ antibody using IF and aligned to Visium slide. Spots with Aβ^+^ staining were labeled as Aβ^+^ ST spots using Loupe Browser. Spot levels (1-3) were assigned based on their distance (level 1: 45 µm, level 2: 145 µm, and level 3: 245 µm) from the site of pathology. (**b**) Co-expression of four modules (M1-M4) for ST spots in control (CT) human MTG based on weighted gene co-expression network analysis (WGCNA) of differentially expressed genes (DEGs) in AD vs control human MTG. (**c**) Co-expression of four modules (M1-M4) in AD human MTG at sites of AD pathology (Aβ, AT8, Aβ/AT8, GFAP/Aβ/AT8, and P2RY12/Aβ/AT8) vs. surrounding ST spots (level 1-3).

**Supplementary Fig. 3.**
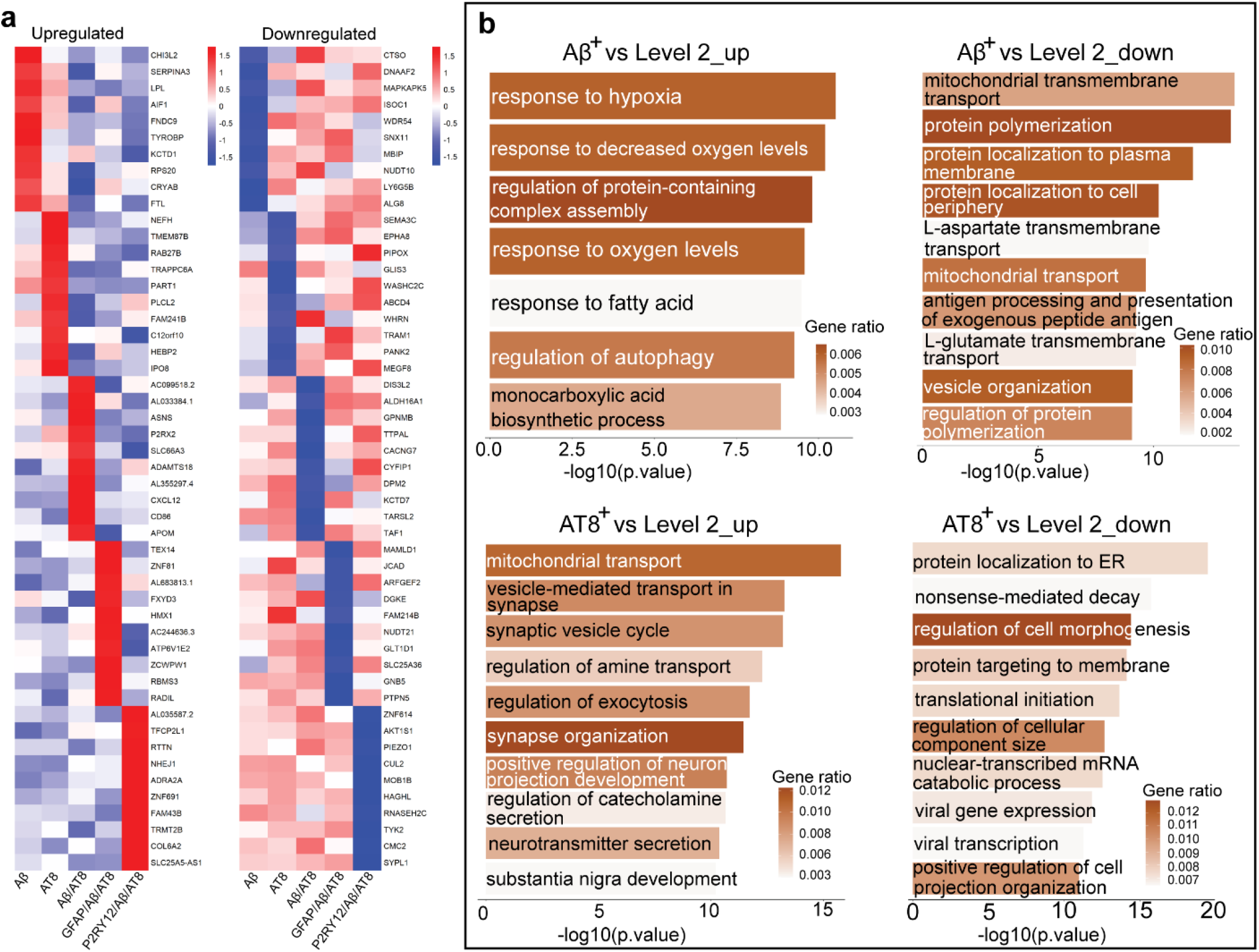
AD-associated gene signatures and pathways at level 2 ST spots. (**a**) Heatmap of Z-scores for upregulated and downregulated differentially expressed genes (DEGs) specific to ST spots localized with and without different AD pathologies (Aβ, AT8, Aβ/AT8, GFAP/Aβ/AT8, and P2RY12/Aβ/AT8). DEGs were identified from five pathological regions vs. surrounding level 2 spots of two AD samples at the pseudo-bulk level. (**b**) Gene ontology (GO) pathway analysis of identified DEGs specific to ST localized with Aβ plaques (top row) or pathological tau, AT8 (bottom row). Pathways and functions associated with both upregulated genes (left) and downregulated genes (right) are shown.

**Supplementary Fig. 4.**
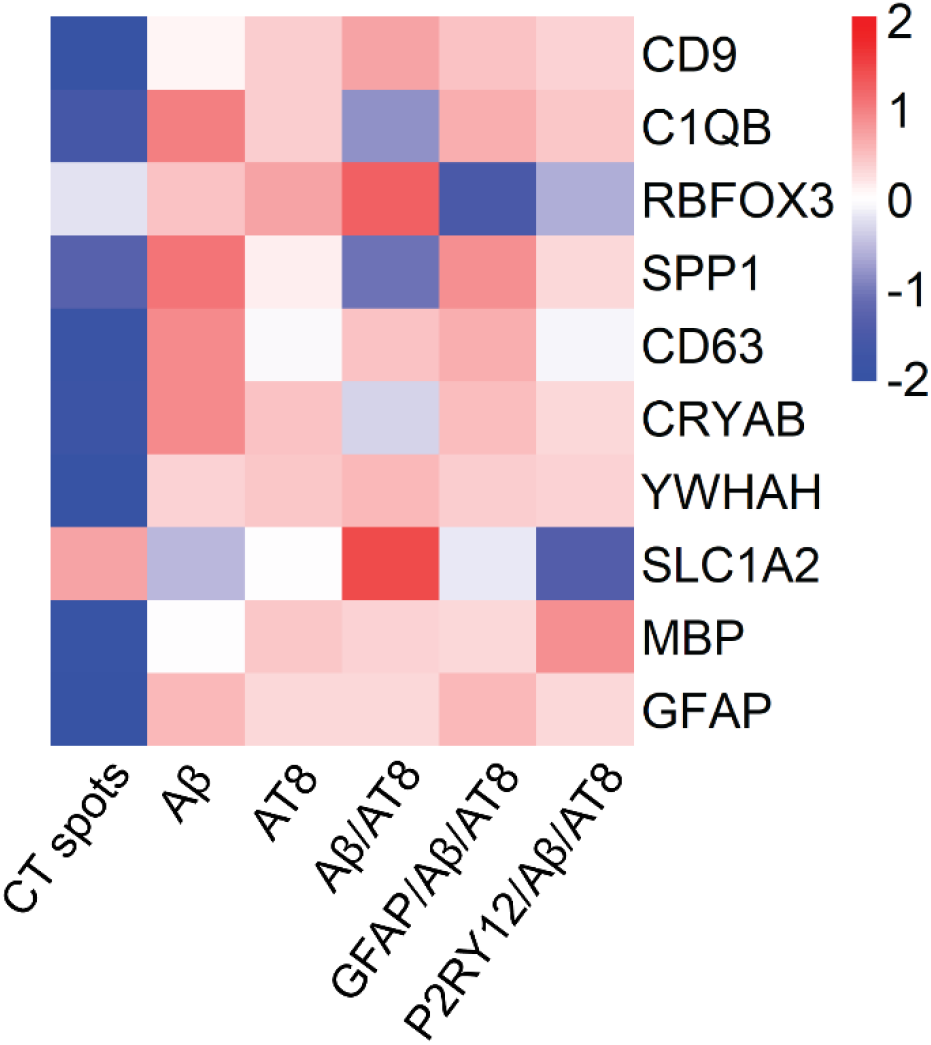
Representative upregulated and downregulated genes associated with AD pathologies. Heatmap of Z-scores for 10 representative differentially expressed genes (DEGs) associated with different AD pathologies (Aβ, AT8, Aβ/AT8, GFAP/Aβ/AT8, and P2RY12/Aβ/AT8) and the control. These 10 DEGs were selected as probes for cell-type specific validation in early AD and control samples using smFISH.

